# Bone marrow AdipoQ-lineage progenitors are a major cellular source of M-CSF that dominates bone marrow macrophage development, osteoclastogenesis and bone mass

**DOI:** 10.1101/2022.07.28.501837

**Authors:** Kazuki Inoue, Yuhan Xia, Yongli Qin, Jean X. Jiang, Matthew B. Greenblatt, Baohong Zhao

## Abstract

M-CSF is a critical growth factor for myeloid lineage cells, including monocytes, macrophages and osteoclasts. Tissue resident macrophages in most organs rely on local M-CSF. However, it is unclear what specific cells in bone marrow produce M-CSF to maintain myeloid homeostasis. Here, we identify bone marrow AdipoQ-lineage progenitors, but not bone marrow mature adipocytes or peripheral adipose tissue, as a major cellular source of M-CSF, with these AdipoQ-lineage progenitors producing M-CSF at levels much higher than those produced by osteoblast lineage cells. Deficiency of M-CSF in bone marrow AdipoQ-lineage progenitors drastically reduces the generation of bone marrow macrophages and osteoclasts, leading to severe osteopetrosis in mice. Furthermore, the postmenopausal osteoporosis in a mouse model can be significantly alleviated by the lack of M-CSF in bone marrow AdipoQ-lineage progenitors. Our findings identify bone marrow AdipoQ-lineage progenitors as a major cellular source of M-CSF in bone marrow and reveal their crucial contribution to bone marrow macrophage development, osteoclastogenesis, bone homeostasis and pathological bone loss.

## Introduction

Macrophage colony-stimulating factor (M-CSF), encoded by the *Csf1* gene, is a growth factor that plays a crucial role in the proliferation, differentiation, survival and function of myeloid lineage cells, including monocytes, macrophages and osteoclasts (1, 2). The global absence of *Csf1* in the *Csf1*^*op*^*/ Csf1*^*op*^ mice, which carry an inactivating mutation in the coding region of *Csf1*, leads to tissue macrophage deficiency, growth retardation and skeletal abnormality. The severe osteopetrosis, shortened long bones and failure of dental eruption in the *Csf1*^*op*^*/ Csf1*^*op*^ mice attest to the essential role of M-CSF for osteoclast generation (3-8).

Osteoclasts are giant multinucleated cells responsible for bone resorption. They are derived from the monocyte/macrophage lineage and are a specialized terminally differentiated macrophage. Osteoclasts play an important role not only in physiological bone development and remodeling, but also actively contribute to musculoskeletal tissue damage and accelerating the progression of postmenopausal osteoporosis and inflammatory arthritis. Osteoclasts and their progenitors express the M-CSF receptor c-Fms. M-CSF is essential for the entire process of osteoclast differentiation, from the generation of osteoclast precursors to the formation and survival of mature osteoclasts (2, 9, 10). M-CSF also acts together with β3 integrin to regulate actin remodeling in osteoclasts to enable osteoclasts to spread, migrate, fuse and form actin rings to facilitate bone resorption (2, 11, 12). These findings underscore an indispensable role for *Csf1* in not only monocyte to macrophage development, but also osteoclast differentiation, function and survival, which directly influences bone mass.

M-CSF is expressed by a variety of cells, such as endothelial cells, myoblasts, epithelial cells and fibroblasts (3, 13-15). This diversity of cellular sources of M-CSF is likely due to the need to support a widely distributed network of tissue resident macrophages, which require M-CSF for their development, function and homeostasis. Under physiologic conditions, circulating M-CSF is mainly produced by vascular endothelial cells (14, 16, 17). The role of circulating M-CSF in tissue macrophage support appears to be highly context dependent. For example, macrophages in kidney and liver are highly dependent on circulating M-CSF. Bone marrow resident macrophages, however, appear to not require circulating M-CSF, indicating the importance of local M-CSF produced in bone marrow (3). Although in vitro studies show that M-CSF is expressed by osteocyte-like cells, an osteoblast cell line, calvarial osteoblastic cells and a cloned bone marrow stromal cell line (9, 18-20), there is no clear evidence supporting whether osteoblasts and osteocytes express M-CSF in vivo and the contributions of osteoblast and osteocyte-derived M-CSF to macrophage and bone homeostasis in vivo is unclear. Therefore, it is largely unknown what specific bone marrow cellular population produces this critical cytokine.

The rapid evolution of scRNAseq technology provides an opportunity to investigate transcriptomics at the individual cell level. In this study, we utilized bone marrow scRNAseq datasets (21-26), and identified a group of unique bone marrow cells featuring AdipoQ expression that highly express *Csf1*. The level of *Csf1* expressed by these bone marrow AdipoQ-lineage progenitor cells is much higher than that produced by osteoblast lineage cells. We further demonstrated the functional importance of the *Csf1* expressed by bone marrow AdipoQ-lineage progenitors in macrophage development, osteoclastogenesis and bone mass maintenance.

## Results

### scRNAseq reveals AdipoQ-lineage progenitors as a main cellular source expressing *Csf1* in bone marrow

Along with the rapid progress of single-cell RNAseq technology, single-cell transcriptomics provides an unprecedented assessment of tissue cellular composition and gene expression profile at individual cell resolution. We took advantage of a recently published dataset (26), which integrated three bone marrow scRNAseq datasets (23-25), and analyzed the expression profiles of non-hematopoietic bone marrow cells. *Adipoq* is found to be exclusively expressed in the cluster MSPC-adipo (mesenchymal progenitor cells-adipo lineage, Fig. 1A, B) as a marker of this cell population. The cellular features of these MSPC-Adipo cells (26) are similar to those described as the adipo-primed mesenchymal progenitors (23), adipocyte progenitors (22), Adipo-CAR (Cxcl12-Abundant Reticular) cells (25), or marrow adipogenic lineage precursors (MALPs) (21) identified in bone marrow. Since we utilized AdipoQ Cre mice to investigate the function of this progenitor population, we used the nomenclature bone marrow AdipoQ-lineage progenitors to designate these cells throughout this study. When screening expression of known genes regulating skeletal homeostasis in the bone marrow, we found that these bone marrow AdipoQ-lineage progenitors (MSPC-adipo cluster) express the highest level of *Csf1* (Fig. 1B, C, D), which was unexpected because the MSPC-osteo (mesenchymal progenitor cells-osteo lineage) cluster and osteoblasts were thought to be the cellular source expressing *Csf1* in bone marrow based on previous studies of in vitro cultured osteoblast cell lines or calvarial osteoblastic cells (9, 18). Further quantitative analysis showed that about 80% of bone marrow AdipoQ-lineage progenitors (MSPC-adipo cluster) express a markedly higher level of *Csf1* than MSPC-osteo cluster cells (Fig. 1C, D). *Csf1* expression was undetectable in osteoblasts (Fig. 1C, D). Interestingly, AdipoQ+ peripheral adipocytes, including both white and brown lipid-laden mature adipocytes, as well as the lipid-laden mature bone marrow adipocytes (BMA), nearly do not express *Csf1* compared to the bone marrow AdipoQ-lineage progenitors that were sorted from the AdipoQ-mTmG mice (Fig. 1E). These results identify AdipoQ-lineage progenitors residing in bone marrow as a new cell type expressing *Csf1*.

**Figure 1.**
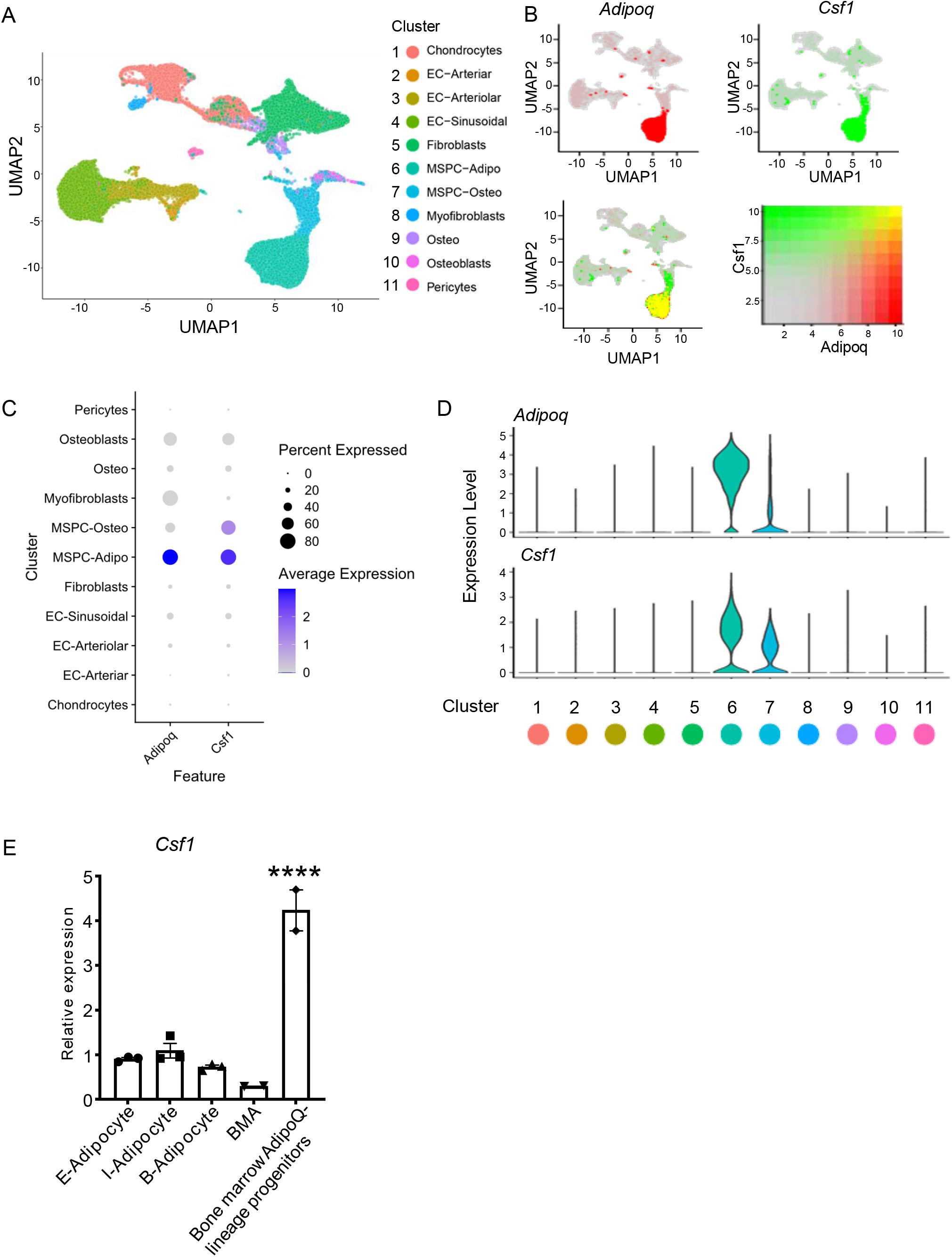
Integrated analysis of the bone marrow niche datasets of scRNAseq shows that AdipoQ+ MSPCs (AdipoQ-lineage progenitors) express high level of Csf1. (**A**) UMAP plot of the integrated analysis of the bone marrow niche datasets of scRNAseq based on (26). EC, endothelial cell; MSPC: mesenchymal progenitor cell. (**B**) UMAP plots of the expression of *Adipoq* (upper left panel), *Csf1* (upper right panel) and the co-expression of these two genes (lower left panel) in bone marrow cells. Lower right panel shows a relative expression scale for each gene. (**C**) Dot plot of *Adipoq* and *Csf1* expression across the listed scRNA-seq clusters. Cell clusters are listed on y-axis. Features are listed along the x-axis. Dot size reflects the percentage of cells in a cluster expressing each gene. Dot color reflects gene expression level as indicated by the legend. (**D**) Violin plots of the expression of *Adipoq* and *Csf1*. (**E**) qPCR analysis of *Csf1* expression in bone marrow AdipoQ-linage progenitors that were sorted from the bone marrow of AdipoQ-mTmG reporter mice, mature bone marrow adipocytes (BMA), mature peripheral white and brown adipocytes. E-adipocyte: mature adipocytes isolated from the epididymal white adipose tissue. I-adipocyte: mature adipocytes isolated from inguinal white adipose tissue. B-adipocyte: Brown adipocytes. n=3 for E-, I- and B-adipocytes from 12-week-old male mice. Two replicates, each with a pooled sample from 12-week-old male mice for BMA (6 mice) and bone marrow AdipoQ-lineage progenitors (3 mice). Error bars: Data are mean ± SEM. ****p < 0.0001.

### AdipoQ Cre-driven *Csf1* conditional knock out (*Csf1*^*ΔAdipoQ*^) mice exhibit osteopetrosis

We next sought to investigate the contribution of the M-CSF produced by bone marrow AdipoQ-lineage progenitors to bone development and homeostasis. We generated *Csf1* conditional knock out (KO) mice, in which *Csf1* is specifically deleted in AdipoQ+ cells by crossing *Csf1*^*flox/flox*^ mice with *AdipoQcre* mice (*Csf1*^*f/f*^*;AdipoQCre;* hereafter referred to as *Csf1*^*ΔAdipoQ*^). Their littermates with a *Csf1*^*f/f*^ genotype were used as the controls. Compared to control mice, *Csf1* expression was reduced by approximately 75% in a total bone marrow stromal cell culture derived from the *Csf1*^*ΔAdipoQ*^ mice (Fig. 2A). Given that bone marrow AdipoQ-lineage progenitors constitute only about 0.08% of bone marrow cells (Suppl. Fig. 1), these results support that bone marrow AdipoQ-lineage progenitors are the major cellular source of M-CSF expression in the bone marrow. *Csf1*^*ΔAdipoQ*^ mice did not display abnormalities in gross appearance, body weight, tooth eruption and long bone length (Fig. 2B, C, Suppl. Fig. 2). In contrast, microcomputed tomographic (µCT) analyses showed that *Csf1*^*ΔAdipoQ*^ mice exhibited a marked osteopetrotic phenotype, indicated by a two-fold increase in trabecular bone mass and marked increases in bone mineral density (BMD), connectivity density (Conn-Dens.), trabecular bone number and decreases in trabecular bone spacing compared to the littermate control mice (Fig. 2D, E). Given that AdipoQ+ cells in peripheral adipose tissue and mature bone marrow adipocytes almost do not express M-CSF, these data indicate that *Csf1* in bone marrow AdipoQ-lineage progenitors plays a key role in the maintenance of bone mass.

**Figure 2.**
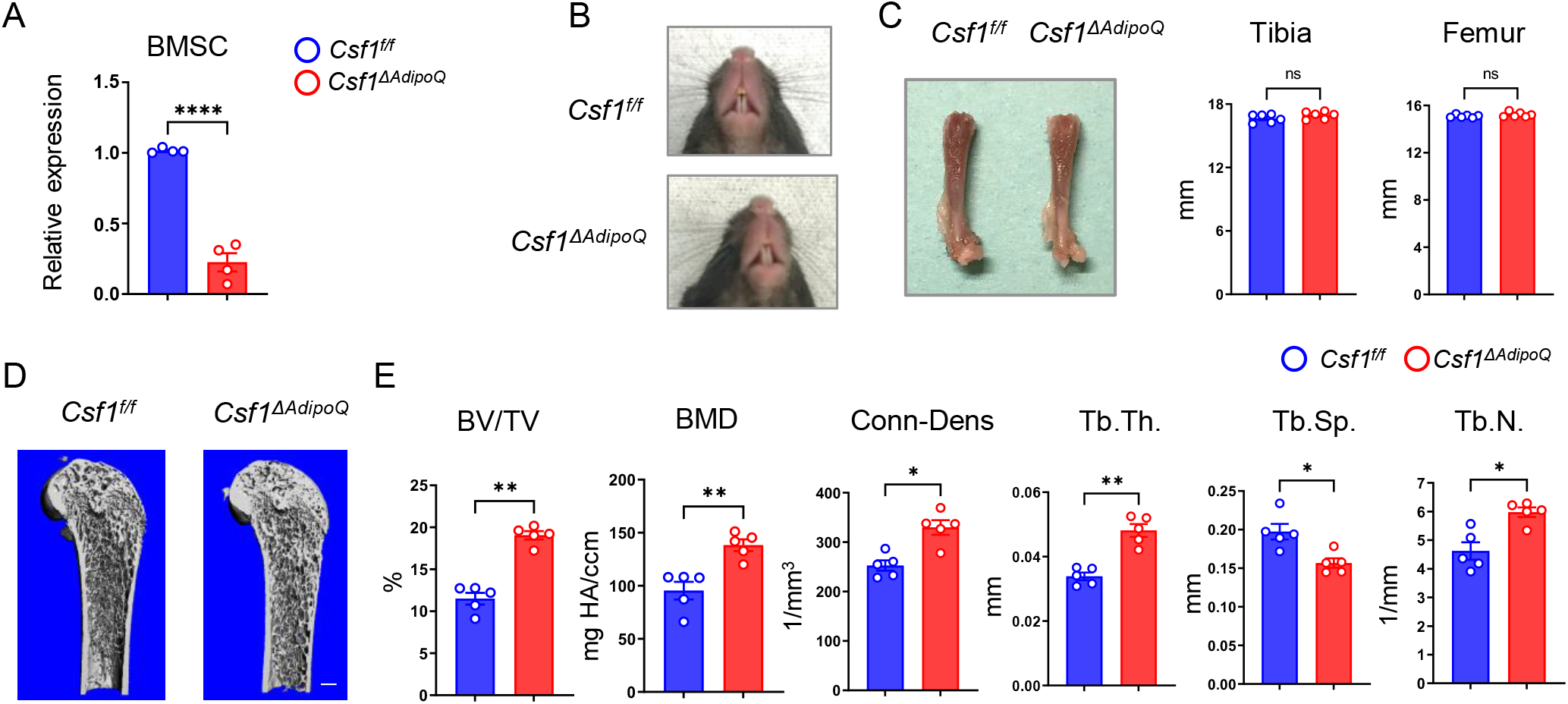
Csf1 deficiency in *Csf1*^*ΔAdipoQ*^ mice increases bone mass. (**A**) *Csf1* expression in BMSCs derived from *Csf* ^*f/f*^ and *Csf1*^*ΔAdipoQ*^ (n = 4/group). (**B**) Gross appearance of the incisors from *Csf* ^*f/f*^ and *Csf1*^*ΔAdipoQ*^ mice. (**C**) Gross appearance of the femur (left panel), and the lengths of femur and tibia from *Csf* ^*f/f*^ and *Csf1*^*ΔAdipoQ*^ mice (right panels) (n = 6/group). (**D**) μCT images and (**E**) bone morphometric analysis of trabecular bone of the distal femurs isolated from 12-week-old male *Csf* ^*f/f*^ and *Csf1*^*ΔAdipoQ*^ mice (n = 5/group). BM, bone marrow; BMSC, bone marrow stromal cell; BV/TV, bone volume per tissue volume; BMD, bone mineral density; Conn-Dens, connectivity density; Tb.Th, trabecular thickness; Tb.Sp, trabecular separation; Tb.N, trabecular number. **A, C, E** *p < 0.05; **p < 0.01; ***p < 0.001; ****p < 0.0001; ns: not statistically significant by two tailed unpaired Student’s t test. Error bars: Data are mean ± SEM. Scale bars: **D**, 500 µm

### Bone marrow macrophages and osteoclasts are suppressed in *Csf1*^*ΔAdipoQ*^ mice

The osteopetrotic phenotype in *Csf1*^*ΔAdipoQ*^ mice implicated a defect in osteoclast function. Indeed, *Csf1* deficiency in AdipoQ+ cells impaired osteoclast formation in vivo evidenced by reduced osteoclast surface area and lower osteoclast numbers (Fig. 3A). On the other hand, mineral apposition rate (MAR) and bone formation rate (BFR) were not affected in *Csf1*^*ΔAdipoQ*^ mice, representing a normal osteoblastic function in these mice (Suppl. Fig. 3). Since osteoclasts are derived from the myeloid macrophage lineage, we examined bone marrow macrophage populations. As expected, CD11b+Ly6C^hi^ monocytes were similar between control and *Csf1*^*ΔAdipoQ*^ mice. CD11b+F4/80+ macrophages were reduced almost half in *Csf1*^*ΔAdipoQ*^ mice (Fig. 3B). This decrease in macrophages reflects the impact from the lack of M-CSF, which is critical for macrophage development, in the *Csf1*^*ΔAdipoQ*^ mice. To further test the importance of the M-CSF produced by bone marrow AdipoQ-lineage progenitors in osteoclastogenesis, we cultured whole bone marrow *ex vivo* without exogenous M-CSF to test whether the AdipoQ-lineage cell-produced M-CSF is sufficient to function together with RANKL to induce osteoclast differentiation. As shown in Fig. 3C, RANKL can induce osteoclast differentiation in the control bone marrow cultures even without addition of M-CSF, but it failed to induce osteoclastogenesis in the *Csf1*^*ΔAdipoQ*^ bone marrow culture, demonstrating that the M-CSF secreted by AdipoQ-lineage progenitors in bone marrow is critical for osteoclastogenesis.

**Figure 3.**
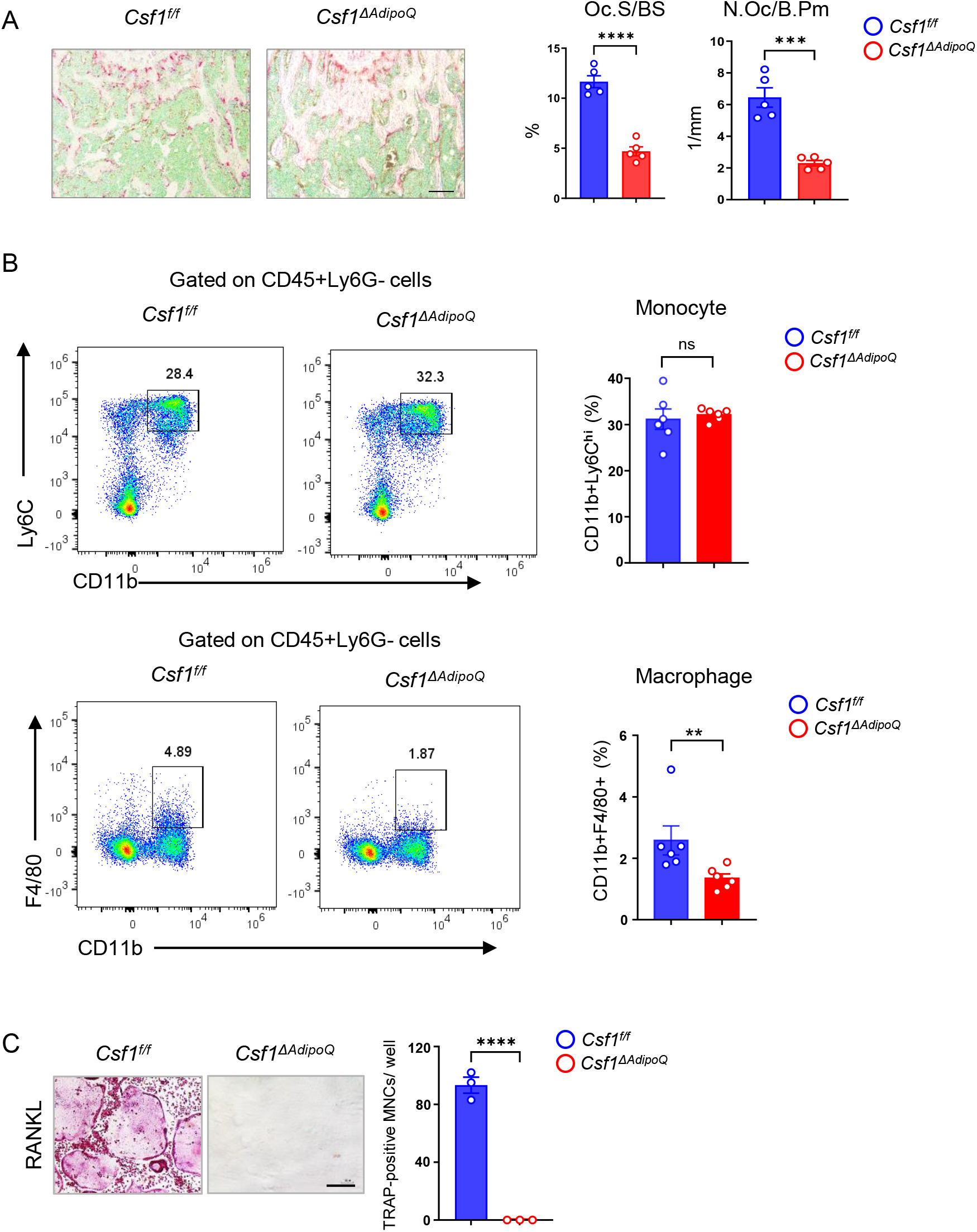
Csf1 deficiency in *Csf1*^*ΔAdipoQ*^ mice suppresses the populations of bone marrow macrophages and osteoclasts. (**A**) TRAP staining and histomorphometric analysis of histological sections obtained from the metaphysis region of distal femurs of 12-week-old male *Csf* ^*f/f*^ and *Csf1*^*ΔAdipoQ*^ mice (n = 5/group). (**B**) Flowcytometry image (left) and quantification (right) of monocytes and macrophages in bone marrow. n=6/group. (**C**) Osteoclast differentiation directly from the cultures of bone marrows harvested from *Csf* ^*f/f*^ and *Csf1*^*ΔAdipoQ*^ mice stimulated with RANKL (40 ng/ml) but without recombinant M-CSF for ten days. TRAP staining (left panel) was performed and the area of TRAP-positive MNCs (≥3 nuclei/cell) per well was calculated (right panel). TRAP-positive cells appear red in the photographs. (n = 3/group). Oc.S/BS, osteoclast surface per bone surface; N.Oc/B.Pm, number of osteoclasts per bone perimeter. **A, B, C** **p < 0.01; ***p < 0.001; ****p < 0.0001; ns: not statistically significant by two tailed unpaired Student’s t test. Error bars: Data are mean ± SEM. Scale bars: **A**, 100 µm; **C**, 200 µm

### *Csf1* deficiency in bone marrow AdipoQ-lineage progenitors does not affect macrophage development in peripheral adipose tissue and spleen

Besides bone marrow, peripheral adiposes contain a large amount of AdipoQ+ mature adipocytes. These cells however do not produce *Csf1* (Fig. 1E). In addition, some organs, such as spleen, have many tissue macrophages but few AdipoQ+ cells. We then asked whether *Csf1* expressed by bone marrow AdipoQ-lineage progenitors affects macrophage population outside bone marrow, such as in peripheral adiposes and spleen. The gross appearance and weight of spleen and peripheral adiposes, such as inguinal and epididymal adiposes, are normal in *Csf1*^*ΔAdipoQ*^ mice (Fig. 4A, B). There is no difference in CD11b+Ly6C^hi^ monocytes in either spleen, inguinal or epididymal adiposes between the control and *Csf1*^*ΔAdipoQ*^ mice, neither does the amount of CD11b+F4/80+ macrophages in these tissues (Fig. 4C). Thus, in contrast to the effects on bone marrow macrophages, *Csf1* deficiency in bone marrow AdipoQ-lineage progenitors does not influence macrophage development in peripheral adipose tissue or spleen.

**Figure 4.**
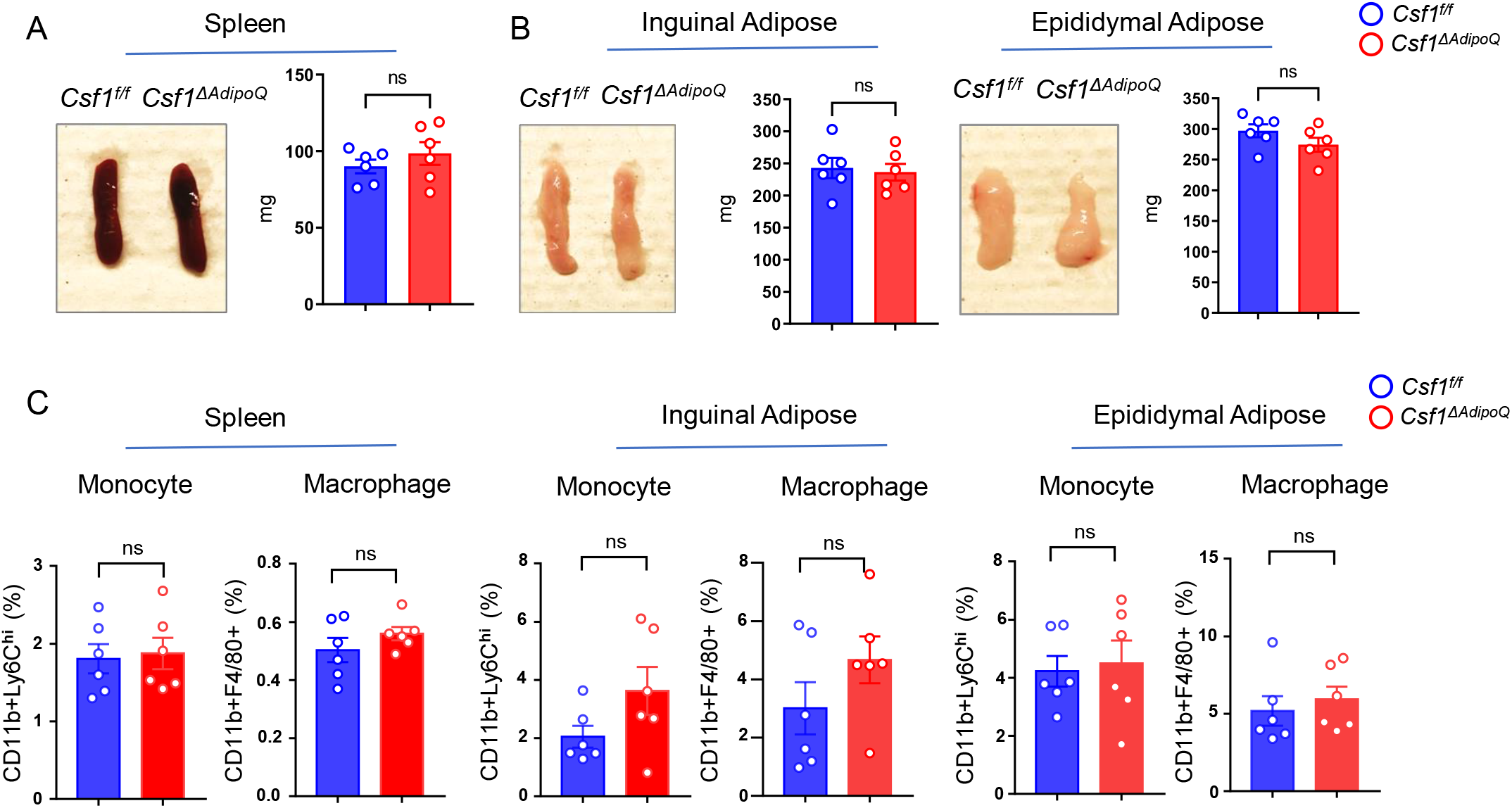
Csf1 deficiency in *Csf1*^*ΔAdipoQ*^ mice does not affect monocyte and macrophage populations in spleen and peripheral adiposes. (**A**) Gross appearance (left panel) and weight (right panel) of the spleen from *Csf* ^*f/f*^ and *Csf1*^*ΔAdipoQ*^ mice (n = 6/group). (**B**) Gross appearance and weight of the inguinal (left panels) and the epididymal (right panels) adipose from *Csf* ^*f/f*^ and *Csf1*^*ΔAdipoQ*^ mice (n = 6/group). (**C**) Flowcytometry quantification of monocytes and macrophages (gated on CD45+Ly6G-cells) in the indicated tissues. n=6/group. **A, B, C** ns: not statistically significant by two tailed unpaired Student’s t test. Error bars: Data are mean ± SEM.

### Lack of *Csf1* in bone marrow AdipoQ-lineage progenitors alleviates estrogen-deficiency induced osteoporosis

We next investigated the significance of the M-CSF secreted by AdipoQ+ cells in pathological bone loss. We developed the ovariectomy (OVX) model in *Csf1*^*ΔAdipoQ*^ mice to study the contribution of AdipoQ+ cell-produced M-CSF to the estrogen-deficiency induced osteoporosis, which mimics postmenopausal bone loss. Uterine weight was measured six weeks after surgery to assess the success of the OVX. Uterine weights were decreased approximately 75% in OVX groups compared with the sham group (Fig. 5A), indicating comparable and effective estrogen depletion in both control and *Csf1*^*ΔAdipoQ*^ mice. μCT analysis showed that OVX significantly reduced bone mass indicated by a decrease in BV/TV, trabecular number, trabecular thickness, Conn-Dens. and an increase of trabecular spacing compared with the sham group in the control mice (Fig. 5B, C). Although OVX also decreased the bone mass of *Csf1*^*ΔAdipoQ*^ mice, the bone loss extent was less in the *Csf1*^*ΔAdipoQ*^ mice than in the control mice (Fig. 5B, C). Furthermore, the bone mass was markedly higher in *Csf1*^*ΔAdipoQ*^ mice than control mice after OVX (Fig. 5B, C, column 4 vs 2), indicating a significant role for the M-CSF secreted by AdipoQ+ cells in estrogen-deficiency induced bone loss. OVX is an osteoclastogenic stimulus. Thus, osteoclast formation as indicated by osteoclast surface and numbers was significantly enhanced by OVX in the control mice (Fig. 5D). Bone histomorphometric analysis further showed significantly greater osteoclast numbers and surface area in the control mice with OVX than the *Csf1*^*ΔAdipoQ*^ mice with OVX (Fig. 5D). Despite an increase in osteoclast surface in *Csf1*^*ΔAdipoQ*^ mice after OVX, osteoclast numbers were not significantly influenced by OVX (Fig. 5D). As bone marrow AdipoQ-lineage progenitors are a key cell population expressing M-CSF, these results indicate that the absence of M-CSF in these cells appears to enable osteoclast linage cells to be resistant to environmental osteoclastogenic stimuli, such as OVX, thereby mitigating pathologic bone loss. These data demonstrate an important role for the M-CSF produced by bone marrow AdipoQ-lineage progenitors in pathologic osteoclastogenesis and bone loss.

**Figure 5.**
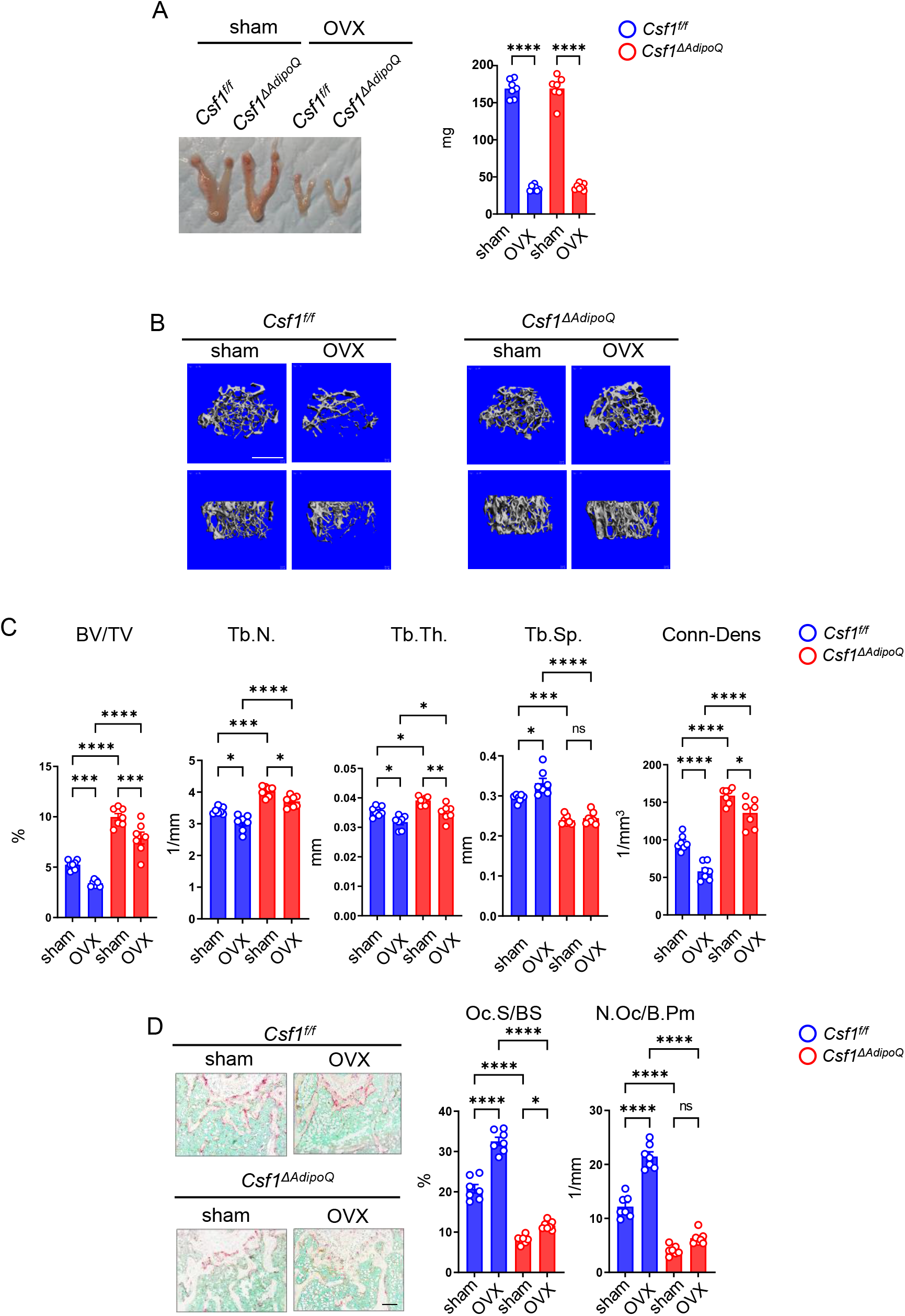
Csf1 deficiency in *Csf1*^*ΔAdipoQ*^ mice protects bone in OVX model.12-week-old female *Csf* ^*f/f*^ and *Csf1*^*ΔAdipoQ*^ mice were subjected to OVX or sham surgery and analyzed 6 weeks after surgery. (**A**) Gross appearance (left panel) and weight (right panel) of uterus, (**B**) μCT images, and (**C**) bone morphometric analysis of trabecular bone of the distal femurs isolated from the *Csf* ^*f/f*^ and *Csf1*^*ΔAdipoQ*^ mice with sham or OVX surgery (n = 7/group). (**D**) TRAP staining (left panels) and histomorphometric analysis (right panels) of histological sections obtained from the metaphysis region of distal femurs isolated from the indicated mice (n = 7/group). BV/TV, bone volume per tissue volume; Tb.N, trabecular number; Tb.Th, trabecular thickness; Tb.Sp, trabecular separation; Conn-Dens., connectivity density; Ct.Th, cortical thickness; Oc.S/BS, osteoclast surface per bone surface; N.Oc/B.Pm, number of osteoclasts per bone perimeter. Data are mean ±SEM. *p < 0.05; **p < 0.01; ***p < 0.001; ****p < 0.0001; n.s., not statistically significant by two-way ANOVA. Scale bars: **B**, 500 µm; **D**, 100 µm

## Discussion

M-CSF as a key cytokine is indispensable for myeloid lineage development and the differentiation and function of osteoclasts, thus playing a crucial role in the physiology and pathology of the immune and skeleton systems. In this study, we identified bone marrow AdipoQ-lineage progenitors as a new cellular source of M-CSF expression, which contributes substantially to the bone marrow macrophage development, physiological bone mass maintenance and pathological bone destruction via controlling osteoclast formation. Bone marrow AdipoQ-lineage progenitors also produce RANKL (27-29), an essential cytokine to induce osteoclast differentiation. These findings collectively highlight the significance of bone marrow AdipoQ-lineage progenitors in the regulation of osteoclastogenesis and bone metabolism, as well as the potential translational implications of appropriately targeting this cell population in treating pathologic bone loss.

AdipoQ is not only highly expressed in the bone marrow AdipoQ-lineage progenitors but also in lipid-laden adipocytes in fat tissues, such as peripheral inguinal and epididymal adipose tissue. However, unlike the wide expression of AdipoQ in lipid-laden adipocytes (BMA) to AdipoQ-lineage progenitors in bone marrow, AdipoQ is exclusively expressed in mature lipid-laden adipocytes but not their progenitors in peripheral adipose tissue. Additionally, unlike bone marrow AdipoQ+ progenitors, these peripheral AdipoQ+ cells nearly do not produce M-CSF. The cells producing RANKL usually appear to simultaneously express M-CSF, such as osteocytes and bone marrow AdipoQ-lineage progenitors (our findings and (21, 27-29)). This is not the case for peripheral AdipoQ+ adipocytes, which express neither RANKL nor M-CSF (our findings and (29)). The unique expression of M-CSF and RANKL by bone marrow AdipoQ-lineage progenitors is a feature that distinguishes this cell population from other adipose depots.

Bone marrow macrophages are reduced by approximately half in the *Csf1*^*ΔAdipoQ*^ mice, but macrophages in other tissues/organs are not significantly affected. These results indicate a strong local regulation of macrophage development by bone marrow AdipoQ-lineage progenitors via M-CSF. It was thought that M-CSF secreted by cells in certain tissues could affect M-CSF levels in other tissues/organs via circulation. However, research using *Csf1*^*op*^*/ Csf1*^*op*^ mice with the restoration of circulating M-CSF shows that some tissues/organs are dependent much more on local than circulating M-CSF, such as bone marrow (3). Our results support this finding and further demonstrate that AdipoQ-lineage progenitors are a crucial cellular source of M-CSF that controls macrophage and osteoclast populations in bone marrow. It is unclear but interesting why macrophage development and osteoclastogenesis in bone marrow is more dependent on local M-CSF. This is presumably a unique mechanism by which bone marrow microenvironment decouples osteoclastogenesis and marrow monocytic development from systemic regulation of tissue resident macrophages, allowing for a marrow homeostasis.

The bone marrow AdipoQ-lineage progenitors only constitute approximately 0.08% of bone marrow cells. However, the deficiency of M-CSF produced by this small AdipoQ+ cell population leads to around a 50% increase in trabecular bone mass and an over 50% reduction bone marrow macrophages. These results together with the scRNAseq data in Fig. 1 demonstrate that AdipoQ-lineage progenitor cells in bone marrow produce substantially more M-CSF than osteoblast lineage cells. Thus, AdipoQ-lineage progenitors are a major cellular source of M-CSF for bone marrow. We also observed that the osteopetrotic phenotype in *Csf1*^*ΔAdipoQ*^ mice is not as strong as *Csf1*^*op*^*/ Csf1*^*op*^ mice. Literature show that M-CSF expression was found in cultures of osteoblastic cells or an osteocyte cell line. Despite scRNAseq analysis showing that osteoblasts do not express *Csf1*, the MSPC-Osteo population does (Fig. 1). The possibility of M-CSF expression by osteocytes also exists, which needs further assessment in vivo. Collectively, the M-CSF produced by MSPC-Osteo cells and potentially osteocytes could partially compensate for the loss of M-CSF from bone marrow AdipoQ-lineage progenitor cells.

M-CSF functions through its receptor c-Fms to exert various biological activities. IL34 was identified later as an additional ligand for c-Fms. Although it also binds to other receptors aside from c-Fms, IL34 has, at least partially, functional redundancy with M-CSF(30, 31). Unlike M-CSF that is broadly and highly expressed by almost every tissue, IL34 is found highly expressed in brain and skin and plays an important role in the differentiation and maintenance of microglia and Langerhans cells(30, 31). The vascular endothelial cells in spleen also produce IL34(32). The results from scRNAseq datasets show that IL34 is expressed in bone marrow, largely by AdipoQ-lineage progenitors but also moderately by pericytes and slightly by MSPC-Osteo cells (26). This indicates that AdipoQ-lineage progenitors are highly likely a major cellular source of IL34 in bone marrow. Spleen has a reservoir of RANK+/CSF-1R+ osteoclast precursors, which can home to bone marrow to differentiate to mature osteoclasts in response to RANKL, IL-34 and/or M-CSF (32). These mechanisms collectively may partially compensate for M-CSF function and alleviate the osteopetrotic and macrophage deficiency phenotype in *Csf1*^*ΔAdipoQ*^ mice.

In addition to its function in development and steady states, M-CSF is a key factor contributing to pathological bone destruction. Broad blockade of M-CSF, however, is not clinically feasible because this may suppress macrophage survival and function in a variety of important tissues and organs. Our results show that lack of M-CSF in bone marrow AdipoQ-lineage progenitors does not affect macrophage populations outside the bone marrow, but can reduce pathological bone loss due to estrogen deficiency. These findings suggest that targeting bone marrow AdipoQ-lineage progenitors-derived M-CSF could be a potential therapeutic strategy to prevent pathological bone destruction.

## Methods

### Mice and analysis of bone phenotype

*Csf1*^*flox/flox*^ mice have been described previously (33). We generated mice with Adipoq+ cell specific deletion of *Csf1* by crossing *Csf1*^*flox/flox*^ mice with the mice with an Adipoq promoter-driven Cre transgene on the C57BL/6 background (AdipoQ-Cre: The Jackson Laboratory, stock No: 028020). We also generated AdipoQ-mTmG reporter mice by crossing the mTmG mice (The Jackson Laboratory, stock No: 007676) with the AdipoQ-Cre mice. Gender- and age-matched *Csf1*^*flox/flox*^*;AdipoQ-Cre* mice (referred to as *Csf1*^*ΔAdipoQ*^) and their littermates with *Csf1*^*flox/flox*^ genotype as controls (referred to as *Csf1*^*f/f*^) were used for experiments. *Csf1*^*flox/flox*^ mice were a gift from Dr. Jean X. Jiang (UT Health San Antonio). Bilateral ovariectomy (OVX) or sham operation (Sham) was performed on 12-week-old female mice. The mice were sacrificed 6 weeks after surgery. Uterine atrophy was first confirmed and then bones were collected for µCT and histological analysis. The mice with the same genotype were randomly allocated to different treatments or procedures. All mouse experiments were approved by Institutional Animal Care and Use Committee of the Hospital for Special Surgery and Weill Cornell Medical College.

µCT analysis was conducted to evaluate bone volume and 3D bone architecture using a Scanco µCT-35 scanner (SCANCO Medical). Mice femora were fixed in 10% buffered formalin and scanned at 6 µm resolution. Proximal femoral trabecular bone parameters or the cortical bone parameters obtained from the midshaft of femurs were analyzed using Scanco software according to the manufacturer’s instructions and the American Society of Bone and Mineral Research (ASBMR) guidelines. Femur bones were decalcified and subjected to sectioning, TRAP staining and histological analysis. For dynamic histomorphometric measures of bone formation, calcein (25 mg/kg, Sigma) was injected into mice intraperitoneally at 7 and 2 days before sacrifice to obtain double labeling of newly formed bones. The non-decalcified femur bones were processed with 5% aqueous potassium hydroxide for 96 hrs, dehydrated and embedded in paraffin (34). 5 μm thick sections were sliced using a microtome. The Osteomeasure software was used for bone histomorphometry using standard procedures according to the program’s instruction.

### Single cell RNAseq (scRNAseq) Analysis

The data from the integrated analysis of the five scRNA-seq datasets of the non-hematopoietic compartment of the murine bone marrow (21-25) were extracted from the Open Science Framework (https://osf.io/ne9vj) (26). We further extracted the clusters of “Chondrocytes”, “EC-Arteriar”, “EC-Arteriolar”, “EC-Sinusoidal”, “Fibroblasts”, “MSPC-Adipo”, “MSPC-Osteo”, “Myofibroblasts”, “Osteo”, “Osteoblasts”, and “Pericytes” from the integrated data and analyzed the data using Seurat (35). UMAP (uniform manifold approximation and projection) was used for dimensionality reduction after PCAs were calculated for integrated datasets. We visualized the simultaneous expression of two genes in a cell using FeaturePlot function in Seurat. Percentage of cells in a cluster expressing a certain gene and the expression level of each gene were visualized by DotPlot function in Seurat. Single cell expression distributions in each cluster were visualized by VlnPlot function in Seurat.

### Cell culture

For osteoclastogenesis directly from bone marrow cultures, bone marrow cells were seeded at a density of 3.125 × 10^5^/cm^2^, and cultured in α-MEM medium with 10% FBS, glutamine (2.4 mM, Thermo Fisher Scientific) and Penicillin–Streptomycin (Thermo Fisher Scientific) for 2 days. The cells were then treated with RANKL (40 ng/ml) without M-CSF for 10 days. Medium were changed every 2 days.

### Reverse transcription and real-time PCR

DNA-free RNA was obtained with the RNeasy MiniKit (Qiagen, Valencia, CA) with DNase treatment, and 1 µg of total RNA was reverse-transcribed with random hexamers and MMLV-Reverse Transcriptase (Thermo Fisher Scientific) according to the manufacturer’s instructions. Real-time PCR was done in triplicate with the QuantStudio 5 Real-time PCR system and Fast SYBR® Green Master Mix (Thermo Fisher Scientific) with 500 nM primers. mRNA amounts were normalized relative to glyceraldehyde-3-phosphate dehydrogenase (GAPDH) mRNA. The primers for real-time PCR were as follows: *Csf1*: 5’-AAAGACAACACCCCCAATGC-3’ and 5’-AGGAGTCTCATGGAAAGTTCGG-3’; *Gapdh*: 5’-ATCAAGAAGGTGGTGAAGCA-3’ and 5’-AGACAACCTGGTCCTCAGTGT-3’.

### Flow cytometry

Bone marrow cells were directly flushed out by quick centrifuge (from 0∼9,400g, approximately 15s at room temperature) after cutting both ends of long bones, then resuspended by PBS (Corning, 21040CV) and filtered through 70 μm cell strainer (FALCON,352350). The cells were spun down at 500g for 5min at 4°C. Epididymal white adipose tissue (EWAT) and Inguinal WAT (IngWAT) were isolated from mice and minced into small pieces by scissors and then digested by the digestion buffer (a-MEM (Gibco,12561056) with 2mg/ml collagenase II (Worthington, LS004176) and 2% BSA (Gemini Bio, 700-100P)) for 40min in a rotary incubator at 250rpm at 37°C. The cells were then filtered through 70 μm cell strainer and spun down at 500g for 5 min at 4°C. One third of spleen was mashed with the plunger end of a syringe through the 70 μm cell strainer. The splenic cells were then spun down at 500g for 5min at 4°C. Red blood cells in cell pellets were lysed by ACK lysis buffer (Gibco, A1049201). Cells were stained with antibodies in 200 μl of the staining buffer (PBS with 0.5% BSA and 2mM EDTA (Invitrogen,15575020)) on ice for 30 min, and then washed by 2 ml of the staining buffer. Cells were resuspended in the staining buffer and analyzed using a FACS Symphony flow cytometer (BD Biosciences). Flowjo V10.7.1 was used for analysis. The antibodies used for flow cytometry included anti-CD45-PerCP/Cyanine5.5 (Biolegend, clone QA17A26,1:200), anti-Ly6G-Brilliant Violet 711 (Biolegend, clone 1A8, 1:200), anti-CD11b-Alexa Fluor 700 (Biolegend, clone M1/70, 1:200), anti-F4/80-APC/Cyanine7(Biolegend, clone BM8, 1:200) and anti-Ly6C-Brilliant Violet 510 (Biolegend, clone HK1.4, 1:200).

### Fluorescence-activated cell sorting (FACS)

Bone marrow cells from 12-week-old male AdipoQ-mTmG reporter mice were harvested as above. The long bones (without periosteum) were then cut into pieces and digested in α-MEM (Gibco,12561056) with 2% FBS (Atlanta Biologicals, S11550), 2mg/ml collagenase II (Worthington, LS004176) and 1mg/ml Dispase II (Gibco, 17105041) for 25 min in a rotary incubator at 250rpm at 37°C. Dnase I (2 units/ml, Sigma,4716728001) was then added. After 5 min, the digested bone solution and bone marrow cells were collected together, filtered through 70 μm cell strainer and spun down at 500g for 5 min at 4°C. Red blood cells in cell pellets were lysed by ACK lysis buffer (Gibco, A1049201). The cells were stained with anti-Lin antibodies, including anti-CD45-APC (Biolegend, clone 30-F11, 1:200), anti-CD31-APC (Biolegend, clone MEC13.3, 1:200) and anti-Ter119-APC (Biolegend, 116212,1:200) for 30 min at 4°C. The AdipoQ+ cells (GFP+) were sorted using a FACS Aria II SORP cell sorter (Becton Dickinson) at Weill Cornell Medical College, with exclusion of DAPI+ (BD, 564907) cells, doublets and Lin+ cells.

### Isolation of peripheral mature adipocytes and bone marrow mature adipocytes

EWAT, IngWAT and brown adipose tissue (BAT) were isolated from mice and minced into small pieces by scissors and then digested by the digestion buffer (a-MEM (Gibco,12561056) with 2mg/ml collagenase II (Worthington, LS004176) and 2% BSA (Gemini Bio, 700-100P)) for 40min in a rotary incubator at 250rpm at 37°C. The cells were then filtered through 70 μm cell strainer and spun down at 500g for 5 min at room temperature. The floating mature lipid-laden adipocytes were collected from the top layer and washed with PBS for 3 times. Bone marrow cells were directly flushed out by quick centrifuge (from 0∼9,400g, approximately 15s at room temperature) after cutting both ends of long bones, then resuspended by PBS (Corning, 21040CV) and filtered through 70 μm cell strainer (FALCON,352350). The cells were spun down at 500g for 5min at room temperature. The floating mature lipid-laden adipocytes were collected from the top layer and washed with PBS for 3 times (29).

### Statistical analysis

Statistical analysis was performed using Graphpad Prism® software. Two-tailed Student’s t test was applied when there were only two groups of samples. In the case of more than two groups of samples, two-way ANOVA was used with more than two conditions. ANOVA analysis was followed by post hoc Bonferroni’s correction for multiple comparisons. p < 0.05 was taken as statistically significant. Data are presented as the mean ± SEM as indicated in the figure legends.

## ACKNOWLEDGEMENTS

We thank Courtney Ng for critical review of the manuscript. We are grateful to the lab members from Dr. Baohong Zhao’s laboratory for their helpful discussions and assistance.

M.B.G. holds a Career Award for Medical Scientists from the Burroughs Welcome Foundation, and a Pershing Square Sohn Prize for Young Investigators in Cancer Research. This work was supported by grants from the National Institutes of Health (MBG, JXJ and BZ), Welch Foundation Grant (AQ-1507 to JXJ) and by support for the Rosensweig Genomics Center at the Hospital for Special Surgery from The Tow Foundation. The content of this manuscript is solely the responsibilities of the authors and does not necessarily represent the official views of the NIH.

## AUTHOR CONTRIBUTIONS

K.I. and Y.X. designed and performed the experiments, analyzed and curated data, prepared figures, and contributed to manuscript preparation. K.I. performed scRNAseq analysis. Y.Q. performed flowcytometry, cell sorting, analyzed and curated data, prepared figures, and contributed to manuscript preparation. J.X.J. provided Csf1 flox mice and contributed to manuscript preparation. M.B.G. provided instruction of experimental designs, discussed data and contributed to manuscript preparation. B.Z. conceived, designed, supervised the project and wrote the manuscript. All authors reviewed, provided input on the manuscript and approved submission.

---

**Suppl Figure 1.**
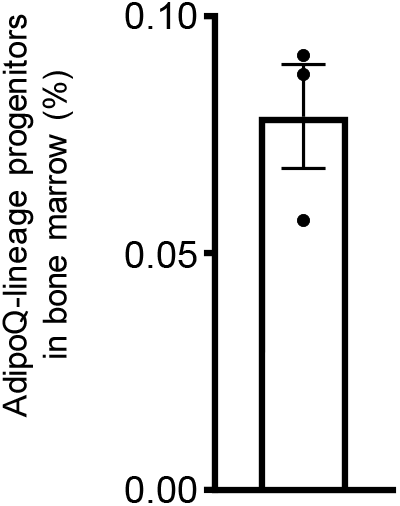
The percentage of AdipoQ-lineage progenitors in bone marrow (without red blood cells) from 12-week-old male mice. n=3. Data are mean ± SEM.

**Suppl Figure 2.**
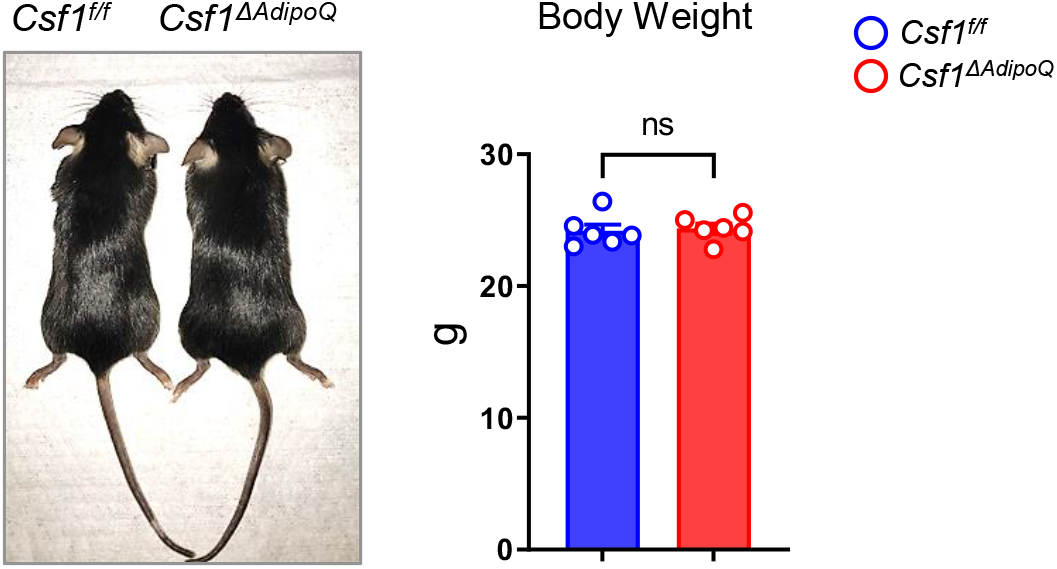
Gross appearance and body weight of 12-week-old male *Csf* ^*f/f*^ and *Csf1*^*ΔAdipoQ*^ mice (n = 6/group). ns: not statistically significant by two tailed unpaired Student’s t test. Data are mean ± SEM.

**Suppl Figure3.**
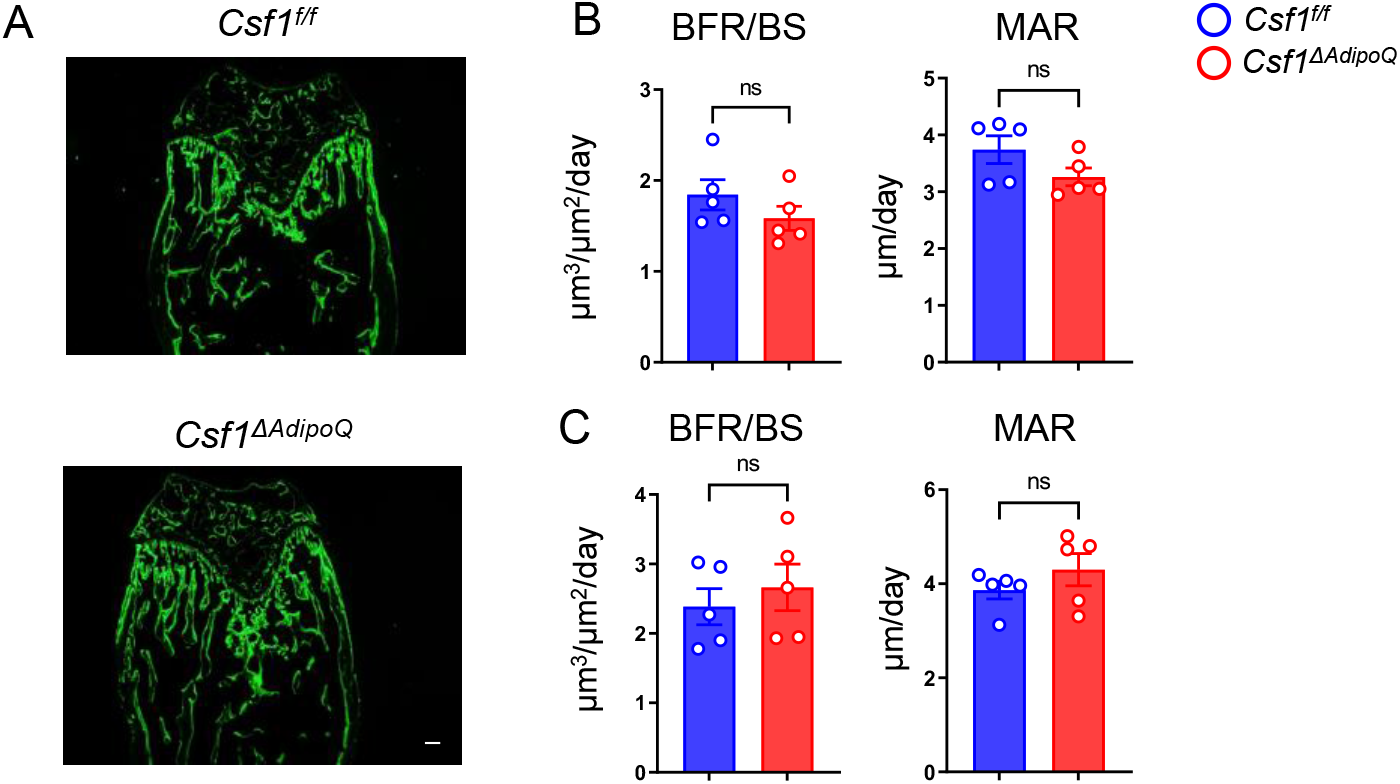
(**A**) Images of calcein double labelling of the femur of 12-week-old male *Csf* ^*f/f*^ and *Csf1*^*ΔAdipoQ*^ mice. Dynamic histomorphometric analysis of trabecular bones (**B**) and cortical bones (**C**) of femurs isolated from 12-week-old male *Csf* ^*f/f*^ and *Csf1*^*ΔAdipoQ*^ mice (n = 5/group). BFR/BS, bone formation rate per bone surface; MAR, mineral apposition rate. Data are mean ±SEM. ns: not statistically significant by two tailed unpaired Student’s t test. Scale bars: **A**, 100 µm

